# Age-Related Impairment of Innate and Adaptive Immune Responses Following Intranasal Herpes Simplex Infection

**DOI:** 10.1101/2022.01.05.475170

**Authors:** Ruchi Srivastava, Anshu Agrawal, Hawa Vahed, Lbachir BenMohamed

**Affiliations:** Laboratory of Cellular and Molecular Immunology, Gavin Herbert Eye Institute, University of California Irvine, School of Medicine, Irvine, CA 92697; Division of Basic and Clinical Immunology, Department of Medicine; University of California Irvine, Irvine, CA 92697-4375, USA; Department of pathology, University of California Irvine, Irvine, CA 92697-4375, USA; Department of Vaccines and Immunotherapies, TechImmune, LLC, University Lab Partners, Irvine, CA 92660-7913; Department of Molecular Biology & Biochemistry; University of California Irvine, School of Medicine, Irvine, CA 92697.; Institute for Immunology; University of California Irvine, School of Medicine, Irvine, CA 92697.

**Author notes:** Corresponding author: Dr. Lbachir BenMohamed, Laboratory of Cellular and Molecular Immunology, Gavin Herbert Institute; Hewitt Hall, Room 232; 843 Health Sciences Rd; Irvine, CA 92697-4390.

**Keywords:** herpes, aging, DCs, CD8^+^ T cells

## Abstract

Immune function declines with age, leading to an increased vulnerability to respiratory viral infections. The mechanisms by which aging negatively impacts the innate and adaptive immune system leading to enhanced susceptibility to infections remain to be fully elucidated. In the present study, we used a mouse model of intranasal infection with herpes simplex virus type 1 (HSV-1), a virus that can enter the lungs through the nasal route causing pneumonia, a serious health concern in the elderly. Following intranasal inoculation of young (6 weeks), adult (36 weeks), and aged (68 weeks) with HSV-1 (KOS strain) we: (*i*) compared the local and systemic innate and adaptive immune response to infection; and (*ii*) correlated the level and type of immune response to protection against HSV-1 infection. Compared to young and adult mice, aged mice displayed: (*i*) increased basal level activation of epithelial cells with a decreased expression of TLR3; (*ii*) increased activation of dendritic cells with increased expression of MHC-1, MHC-II and CD80/86; and (*iii*) decreased production of type-I interferons upon stimulation; (*iv*) a delay in cytokines and chemokines production in the lungs; and (*v*) an impairment in function (CD107 and IFN-γ production) of HSV-specific CD8^+^ T cells. These impairment in innate and adaptive immune responses in aged mice following intranasal HSV-1 inoculation was associated with symptomatic herpes infection. The findings suggest an age-related impairment of both innate and adaptive immune responses which may exacerbate herpes infection and disease in the elderly.

**IMPORTANCE:** Immune function declines with age, leading to an increased vulnerability of the elderly to respiratory viral infections. The mechanisms by which aging negatively impacts the innate and adaptive immune system leading to enhanced susceptibility to respiratory infections remain to be fully elucidated. The present study showed that, compared to young and adult mice, aged mice displayed increased basal level activation of epithelial cells with a decreased expression of TLR3 increased activation of dendritic cells associated with an impairment of HSV-specific CD8^+^ T cell responses. These immune dysregulations were associated with to symptomatic herpes infection. The findings suggest an age-related impairment of both innate and adaptive immune responses which may exacerbate herpes infection and disease in the elderly.

## INTRODUCTION

Herpes simplex virus type 1 (HSV‐1) causes a variety of infections that involve mucocutaneous surfaces, the central nervous system and, occasionally, visceral organs such as the lungs (1), (2). Seventy percent of the human population infected with HSV are adults above the age of 65.Infection of the respiratory tract with HSV causes herpes simplex pneumonia (HSP). Immune function declines with age, making elderly people more susceptible to pulmonary viral infections.

The virus has been reported to be associated with pulmonary disease since 1949 (3). Pulmonary innate immune responses depend upon a highly regulated multicellular network to defend an enormous surface area of interaction with the external world. Local disruption of these responses renders the host susceptible to respiratory infections and subsequent systemic spread of infection. Studies within the last decade determined that airway epithelial cells, dendritic cells (DCs) and macrophages are key participants in this innate immune network. DCs are the major antigen presenting cells which can sense and respond to pathogens and activate T cells in the lungs.

The increased morbidity and mortality reported in elderly populations are due to several factors that include dysfunctions in the senescent immune system. Peripheral immune responses are virtually constant in young individuals but are invariably reduced in aged mice and humans. Several *in vitro* and *in vivo* studies have pointed to the type I interferon (IFN) response as a critical pathway involved in the early immune response to infection (4). Multiple pattern recognition receptors, such as the toll-like receptor 3 (TLR-3) have also been demonstrated to be involved in controlling viral replication in murine models of infection (5).

Since the elderly population is increasing worldwide, a better understanding of the changes that the immune system undergoes with the aging process is becoming a key factor in the development of new therapeutic strategies. The decline in the immune function of the elderly, places them at a high-risk category for clinical herpes virus reactivation and/or severe disease. Thus, the development of novel therapeutic targets is urgently needed. Additionally, associations between innate and adaptive immune responses of the elderly population have not been fully examined in herpes simplex infection and disease. Thus, in this report, we explored the age-related immune cell responses (in terms of magnitude, functional capacity, and repertoire diversity) in young mice (6 weeks of age), adult mice (36 weeks of age), and aged mice (68 weeks of age) following intranasal herpes simplex virus type 1 (HSV-1) infection. After infection with herpes simplex virus, production of interferon alpha (IFN-α) was reduced leading to a significantly reduced T cell response in old compared to young mice. More importantly, control of virus replication was profoundly impaired in aged mice compared to young mice. HSV-infection expanded the total CD8^+^ T cell and CD4^+^ T cells, however, the overall immune response decreases with age. Our findings demonstrated that: (**1**) aged mice are unable to mount a protective immune response against pulmonary HSV infection while the young mice displayed robust immunity; (**2**) The expression of pathogen recognition receptors (PRRs) such as TLR3 was reduced in aged mice; (**3**) There was reduced production of type I Interferon; (**4**) The functions of innate immune cells including epithelial and dendritic cells were compromised in aged mice. The expression of MHC I and II and the co-stimulatory molecules are higher in aged DCs indicating an activated state; and (**5**) The CD8^+^ T cell responses against HSV-1 were reduced in the aged mice. In summary, our results demonstrated an impaired immune responses to herpes infection in aged mice compared to young mice, further supporting immune-senescence in the elderly.

## MATERIALS and METHODS

### Mice

Six week of age (young), 36 week of age (adult) and 68 week of age (aged) female C57BL/6 (B6) mice were purchased from The Jackson Laboratory (Bar Harbor, ME). Animal studies conformed to the Guide for the Care and Use of Laboratory Animals published by the US National Institute of Health (IACUC protocol # AUP 19-111).

### Virus production

Herpes simplex virus type 1 (HSV-1, strain KOS) was grown and titrated on rabbit skin (RS) cells as previously described (6-9).

### Intranasal infection

6-, 36- and 68-week old mice were anesthetized with a mixture of ketamine and xylazine and intranasally infected with 50 μl of PBS containing 1×10^6^ pfu of HSV-1 (KOS strain).

### Nasal swab collection

Nasal swabs were collected on day 2 and day 6 post-infection by pipetting 30 μL of phosphate-buffered saline (PBS) in and out of the nasal openings three times. Swabs were then frozen at −80°C.

### Viral titer assay

Nasal washes and lung tissue were analyzed for viral titers by plaque assays. Positive controls were run with every assay with previously titrated laboratory stocks of HSV-1.

Briefly, Vero cells were grown in an α-modified Eagle’s medium (ThermoFisher Scientific, Waltham, MA) supplemented with 5% fetal bovine serum and 1% penicillin-streptomycin, and L-glutamine (ThermoFisher Scientific). For plaque assays, Vero cells were grown to confluence in 24-well plates. Nasal wash samples were added to monolayers. Infected monolayers were incubated for 1 hour at 37°C and were rocked every 15 minutes for viral absorption. Infected monolayers were overlaid with media containing carboxy-methyl cellulose. Infection was allowed to occur for 72 hours at 37°C. Monolayers were then fixed and stained with crystal violet. Subsequently, the viral plaques were counted under a light microscope.

### Tissue Harvesting and lung cell isolation

Lungs were harvested (n = 5 per time point per experiment) on days 2 and 6 post-infection. Lungs were exposed by opening the chest cavity and rinsed with cold 1× PBS, through the right heart ventricle. Mice lung tissues were harvested, minced, and digested in 5 mg/ml collagenase for 45 minutes, before filtering through a 70 μm cell strainer. Subsequently, cells were spun down and diluted to 1 × 10^6^ viable cells per ml in RPMI media with 10% (v/v) FBS. Viability was determined by Trypan blue staining.

### Flow cytometry

Single cell suspensions from the lungs, were prepared for flow cytometric analysis. The following antibodies were used: anti-mouse CD8 PerCP (BD Biosciences, San Jose, CA), anti-mouse CD11b FITC (BD Biosciences), anti-mouse CD103 APC (BD Biosciences) anti-mouse CD11c (BD Biosciences) anti-mouse CD45 APC-cy7 (BioLegend, San Diego, CA), anti-mouse TLR3 (BD Biosciences), anti-mouse MHCI (BD Biosciences), MHCII (eBioscience), CD4 Percp, CD107^a^ FITC, CD107^b^ FITC (BD Biosciences) and anti-mouse IFN-γ PE-cy7 (clone XMG1.2, BioLegend). For surface staining, mAbs were added against various cell markers to a total of 1 ×10^6^ cells in phosphate-buffered saline (PBS) containing 1% FBS and 0.1% Sodium azide (fluorescence-activated cell sorter [FACS] buffer) and left for 45 minutes at 4°C. For intracellular staining, cells were first treated with cytofix/cytoperm (BD Biosciences) for 30 minutes. Upon washing with Perm/Wash buffer, mAbs were added to the cells and incubated for 45 minutes on ice in the dark. Cells were subsequently washed with Perm/Wash and FACS buffer and fixed in PBS containing 2% paraformaldehyde (Sigma-Aldrich, St. Louis, MO).

For the measurement of CD107^a/b^ and IFN-γ, 1×10^6^ cells were first transferred into 96-well flat bottom plate in the presence of BD GolgiStop (10 μg/ml) for 6 hours at 37°C. Phytohemagglutinin (PHA) (5 μg/ml) (Sigma-Aldrich) was used as positive control. At the end of the incubation period, the cells were transferred to a 96-well round bottom plate and washed once with FACS buffer. Surface and intracellular staining were performed as described previously. A total of 100,000 events were acquired by the LSRII (Becton Dickinson, Mountain View, CA) and analyzed with the FlowJo software (TreeStar, Ashland, OR).

### Cytokine Analysis

Lung cells collected on day 2 and day 6 post-infection was cultured overnight and supernatants collected were assayed for cytokines and chemokines using the multiplex magnetic bead-based kit (ThermoFisher Scientific).

### Statistical analysis

We examined the distribution of each immunological parameter (6). In the case of two group comparisons, we considered the use of the parametric two-sample Student’s *t*-test or non-parametric Wilcoxon rank sum test. In addition, for paired comparisons involving multiple peptides, we have adjusted for multiple comparisons using the Bonferroni procedure. In the specific case of three groups (comparing two subgroups with a baseline subgroup) we used the General Linear Model procedure and compared the least squares means using the Dunnett procedure for multiple comparisons. Flow cytometry data were analyzed with FlowJo software (TreeStar). SAS^®^ v.9.4 (Statistical Analysis System, Cary, NC) was used for analysis. Graphs were prepared with GraphPad Prism software (San Diego, CA). Data are expressed as the mean + SD. Results were statistically significant at *p* < 0.05.

## RESULTS

### 1. Decreased TLR-3 expression and increased activation of airway epithelial cells (AECs) in aged mice at homeostasis

Airway epithelial cells (AEC) are the first cells to sense and respond to infections (10). Viral replication takes place primarily in the epithelium. Herpes infection signals via TLR3 to elicit antiviral innate immune responses in host cells (11). TLR3 is a pattern recognition receptor that recognizes viral double stranded RNA produced during viral replication including HSV-1(12). Herein, we studied the role of AEC responses in the innate immune defense against intranasal infection with HSV-1. Six-, 36- and 68-week old mice were infected intranasally with 1 × 10^6^ pfu of HSV-1 strain (KOS). Two days post infection, mice were euthanized and single cell suspension from the lungs were obtained after collagenase treatment. CD45^−^EPCAM^+^ AECs were analyzed for the expression of TLR3, ICAM-1 (CD54) and MHC-I via flow cytometry (**Fig. 1A**). AECs from aged mice displayed significantly reduced expression of TLR3 at homeostasis as compared to adult and young mice, indicating a reduced capacity to sense HSV-1 infection (**Fig. 1B**). In contrast, the expression of ICAM-1 and MHC-I in AECs was higher in aged mice at homeostasis. Since these are markers of activated epithelium, our results suggest that the airway epithelium is inherently activated at baseline in the absence of infection (**Fig. 1C** and **1D**). Upon infection, the expression of ICAM-1 and MHC-I was upregulated in young and adult mice, but there was no change in aged mice suggesting that aged AECs were not responding to HSV-1 infection (**Fig. 1C** and **1D**). Altogether, these results indicate that AECs from aged mice are impaired in their capacity to sense, respond to infections, and display an increased basal level of epithelial activation.

**Figure 1:**
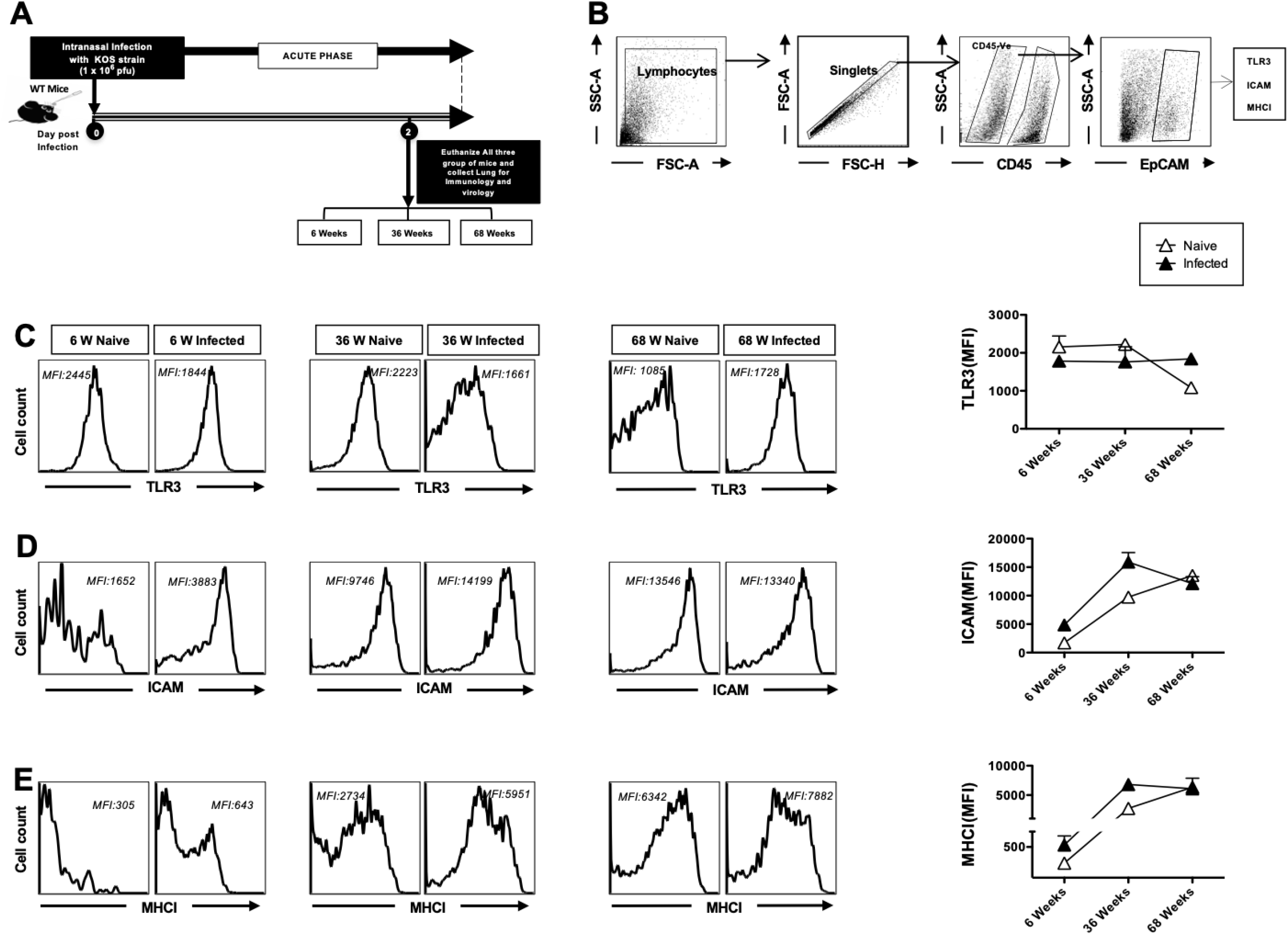
Decreased TLR3 expression and increased activation of airway epithelial cells (AECs) in aged mice at homeostasis: 6-week, 36-week and 68-week-old mice were infected intranasally with 1 × 10^6^ pfu of HSV-1 strain. Mice were euthanized 2 days post infection and a single cell suspension from lungs were obtained after collagenase treatment. The lung cells were stained for epithelial cell markers, TLR-3 and activation markers and then analyzed by FACS. (**A**) Timeline of infection and immunological analyses. (**B**) Gating strategy used to characterize lung derived cells. Lymphocytes were identified by a forward scatter (FSC) and side scatter (SSC) gate. Singlets were selected by plotting forward scatter area (FSC-A) vs. forward scatter height (FSC-H). CD45 negative cells were then gated. The epithelial cell population in CD45-gated cells was defined by the EpCAM^+^ cells. (**C**) Representative histograms and average mean fluorescence intensity (MFI) of TLR-3 expression on the surface of lung epithelial cells (**D**) ICAM expression on epithelial cells (**E**) MHCI expression on lung epithelial cells of 6-week, 36-week and 68-week-old mice at 2 days post-infection with HSV-1 (solid black) or mock-infection with DMEM (solid white). Data is representative of two separate experiments.

### 2. The expression of TLR3 on DCs and macrophages in the lung decreases with age at homeostasis

Next, we examined the expression of TLR3 on DCs and macrophages since these cells also get activated via TLRs (**Fig. 2A**). The single cell lung suspension prepared above (**Fig. 1**) was also utilized for these experiments. To elucidate the consequences of aging on TLR expression on various populations of DCs, we utilized multicolor flow cytometry and intracellular cytokine staining in lung samples from young, adult, and old mice (**Fig. 2B)**; lymphocytes were identified by a forward scatter (FSC) and side scatter (SSC) gate. Singlets were selected by plotting forward scatter area (FSC-A) versus the forward scatter height (FSC-H). Subsequently, CD45 positive cells were gated by the expression of CD45. The myeloid cell population in CD45^+^ gated cells was defined by the CD11b^+^ CD11c^+^ cells (mDC). Similarly, CD11b^+^ subset was defined as macrophages and CD11c^+^ cells as lymphoid cell subsets. TLR-3 expression was analyzed on the surface of myeloid cell subset (CD11b^+^CD11c^+^), macrophages (CD11b^+^CD11c^−^) and lymphoid dendritic cells (CD11c^+^) from 6-, 36- and 68-week-old mice 2 days post-infection. We found that the TLR-3 expression was decreased at baseline in 68-week-old mice as compared to adult and young mice (**Fig. 2C, 2D** and **2E)**. The difference between the groups was not significant after infection. Decreased TLR3 expression at baseline in DCs and macrophages from aged mice is indicative of their compromised capacity to sense and respond to TLR3 ligands including HSV-1, rendering them more susceptible to these viral infections.

**Figure 2:**
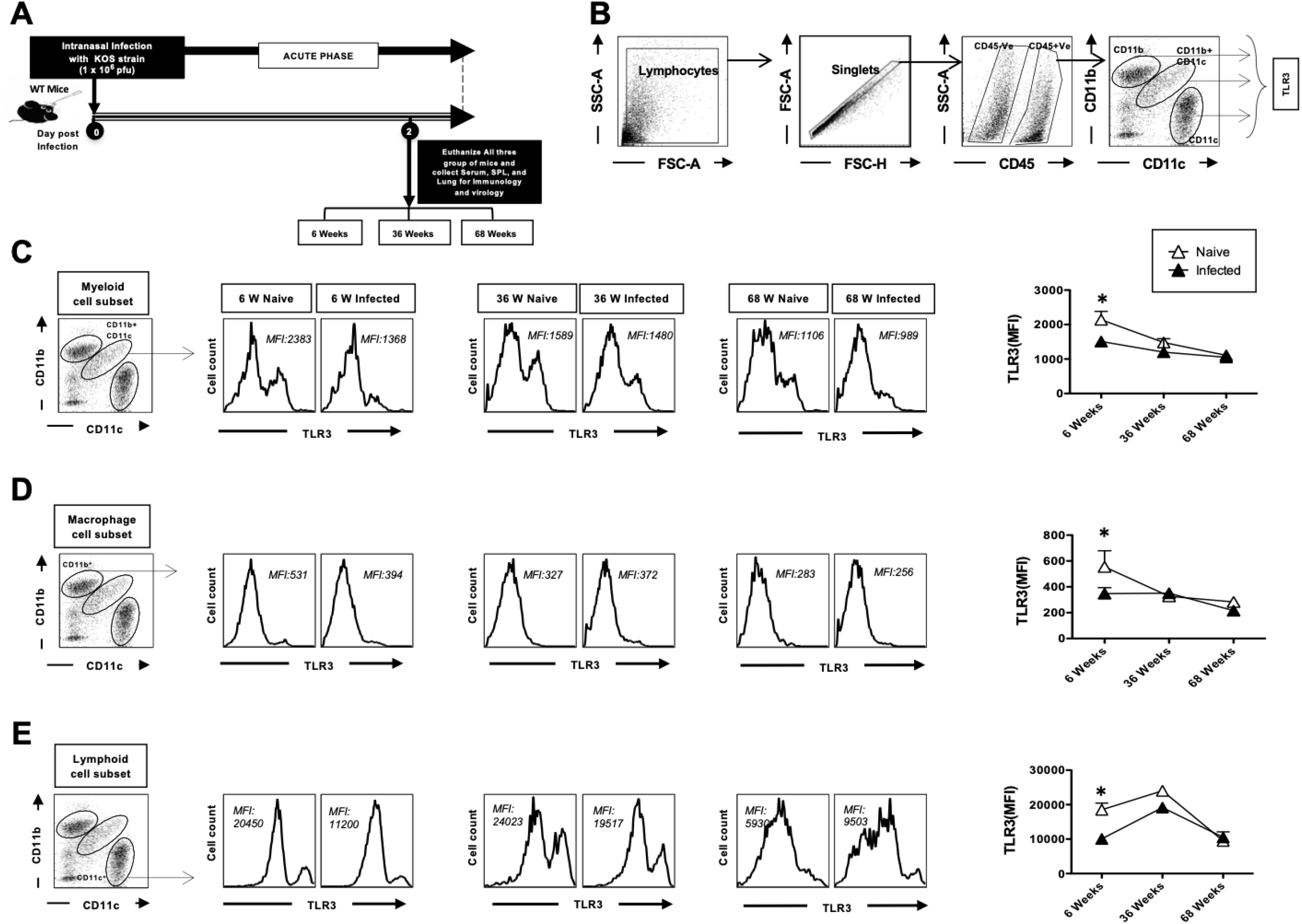
The expression of TLR3 on DCs and macrophages in the lung decreases with age at homeostasis. 6-week, 36-week and 68-week-old mice were infected intranasally with 1 × 10^6^ pfu of HSV-1 strain. Two days post infection mice were euthanized and single cell suspension from lungs were obtained after collagenase treatment. The lung cells were stained for dendritic cell markers and activation markers and then analyzed by FACS. (**A**) Timeline of infection and immunological analyses. (**B**) Gating strategy used to characterize lung derived cells. Lymphocytes were identified by a forward scatter (FSC) and side scatter (SSC) gate. Singlets were selected by plotting forward scatter area (FSC-A) vs. forward scatter height (FSC-H). CD45 positive cells were then gated by the expression of CD45. The myeloid cell population in CD45^+^ gated cells was defined by the CD11b^+^ CD11c^+^ cells. Similarly, CD11b^+^ subset was defined as macrophages and CD11c^+^ cells as lymphoid cell subsets. Representative histograms and average mean fluorescence intensity (MFI) of TLR-3 expression on the surface of (**C**) myeloid cell subset (CD11b^+^CD11c^+^), (**D**) macrophages (CD11b^+^CD11c^−^) and (**E**) dendritic cells (CD11c^+^) from 6-week, 36-week and 68-week-old mice at 2 days post-infection with HSV-1 (solid black) or mock-infection with DMEM (solid white). Data is representative of two separate experiments.

### 3. The upregulation of MHC-I after herpes infection is impaired with age

Since DCs and macrophages are antigen-presenting cells, we examined the up-regulation of MHC class I and MHC class II cell surface markers on these cells. The upregulation of MHC-I on cells is required for priming CD8^+^ T cell responses. The surface expression levels of MHC-I on lung mDCs, macrophages and lymphoid DCs collected from young, adult, and aged mice. Only aged mice showed upregulated levels of MHC-1 on lung mDCs, macrophages, and lymphoid DCs at baseline without infection. Two days post infection, only mDCs and macrophages from young mice displayed significant up-regulation of MHC-I, while no significant change was seen in the adult versus the aged mice (**Fig. 3A**). However, lung macrophages MHC class I expression was similar between the young, adult, and aged HSV-1 infected mice groups. Despite that lung macrophages from adult mice appeared to up-regulate MHC class I quicker than macrophages from young animals (**Fig. 3B**). Similar MHC-I up-regulation patterns on lung lymphoid DCs from young, adult and aged mice were detected (**Fig. 3C**).

**Figure 3:**
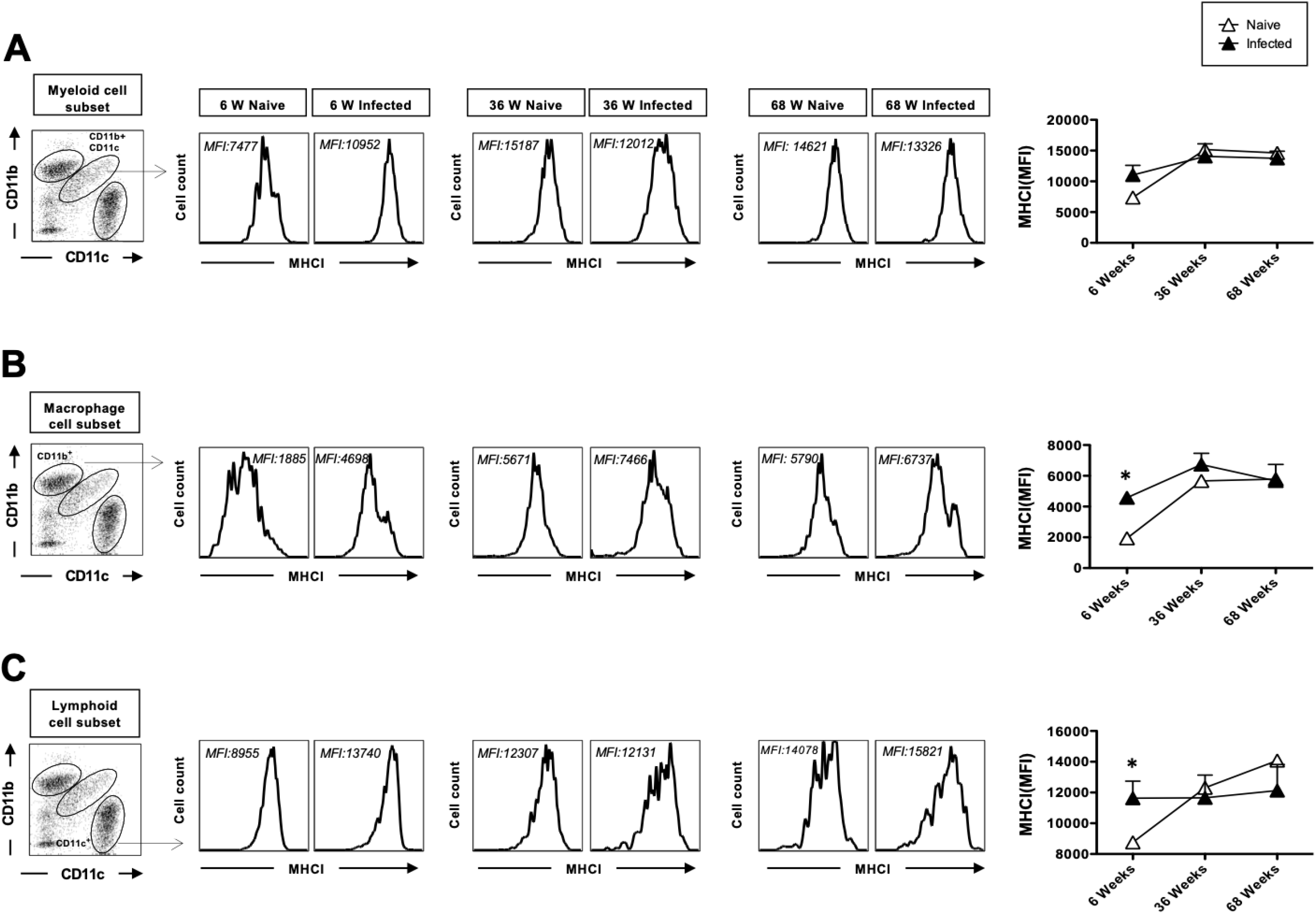
The upregulation of MHC-I after herpes infection is impaired with age. 6-week, 36-week and 68-week-old mice were infected intranasally with 1 × 10^6^ pfu of HSV-1 strain. 2 days post infection mice were euthanized and single cell suspension from lungs were obtained after collagenase treatment. The lung cells were stained for dendritic cell markers and activation markers and then analyzed by FACS. CD45 positive cells were then gated by the expression of CD45. The myeloid cell population in CD45^+^ gated cells was defined by the CD11b^+^ CD11c^+^ cells. Similarly, CD11b^+^ subset was defined as macrophages and CD11c^+^ cells as lymphoid cell subsets. Representative histograms of MHC-I expression mean fluorescence intensity (MFI) on the surface of (**A**) myeloid cell subset (CD11b^+^CD11c^+^), (**B**) macrophages (CD11b^+^CD11c^−^) and (**C**) dendritic cells (CD11c^+^) from 6-week, 36 week and 68-week-old mice at 2 days post-infection with HSV-1 (solid black) or mock-infection with DMEM (solid white). Data is representative of two separate experiments.

### 4. The upregulation of MHC-II after herpes infection is impaired with age

We next examined the effect on MHC-II expression on DCs and macrophages. DCs and macrophages upon activation upregulate MHC-II expression helping them to present antigens to CD4 T cells and therefore activating them. The surface expression levels of MHCII on lung mDCs, macrophages and lymphoid DCs collected from aged mice was upregulated at baseline without infection compared to young and adult mice. The cell surface expression of MHCII on lung mDCs was significantly up-regulated 2 days post-infection in young mice, where there was no increase observed in in adult and aged mice (**Fig. 4A**). MHC class II expression was similar on lung macrophages although adult mice did display a higher MHC-II expression at baseline and post-infection as compared to young and old mice (**Fig. 4B**). Lymphoid DCs displayed a similar pattern of MHC-II expression as the mDCs (**Fig. 4C**).

**Figure 4:**
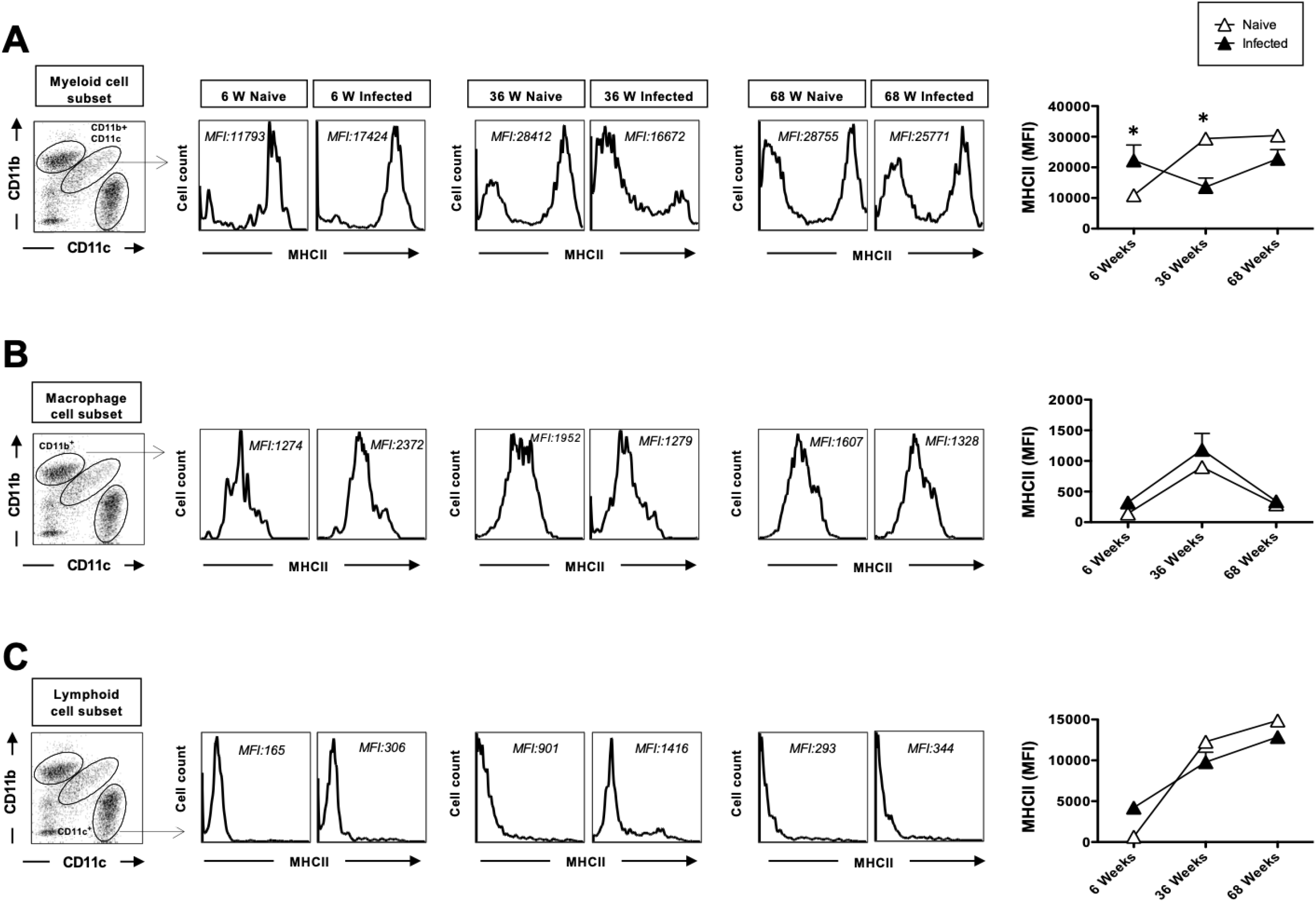
The upregulation of MHC-II after herpes infection is impaired with age. 6-week, 36-week and 68-week-old mice were infected intranasally with 1 × 10^6^ pfu of HSV-1 strain. Two days post infection mice were euthanized and single cell suspension from lungs were obtained after collagenase treatment. The lung cells were stained for dendritic cell markers and activation markers and then analyzed by FACS. CD45 positive cells were then gated by the expression of CD45. The myeloid cell population in CD45^+^ gated cells was defined by the CD11b^+^ CD11c^+^ cells. Similarly, CD11b^+^ subset was defined as macrophages and CD11c^+^ cells as lymphoid cell subsets. Representative histograms mean fluorescence intensity (MFI) of MHC-II expression on the surface of (**A**) myeloid cell subset (CD11b^+^CD11c^+^), (**B**) macrophages (CD11b^+^CD11c^−^) and (**C**) dendritic cells (CD11c^+^) from 6-week, 36 week and 68-week-old mice at 2 days post-infection with HSV-1 (solid black) or mock-infection with DMEM (solid white). Data is representative of two separate experiments.

### 5. Aged mice display reduced secretion of protective cytokines in the lung after acute HSV infection

In this study, we determined the influence of age on the production of interferon (IFN) alpha, IL-1beta, RANTES, IL-22 and IFN-γ. Cytokines play an important role in establishing an antiviral state as the nonspecific first line of defense in viral infections. Changes due to aging occur among a wide variety of different immune parameters, such as the induction of cell surface activation markers, secretion of cytokines, and proliferative capacity. We utilized a multiplex detection to quantitate the levels of inflammatory mediators in the lungs of 6-, 36- and 68-week-old mice on day 2 post infection. The interferon (IFN) alpha, IL-1beta, RANTES, IL-22 and IFN-γ cytokines and chemokines were measured in the culture supernatants after *in vitro* culture of the cells for 24 hours. We detected that IFN-α and IL-1β expression was significantly increased in young mice after herpes infection while there was no change observed in adult and old mice suggesting a reduced response to infection with age (**Fig. 5**). Both IFN-α and IL-1β play an important role in the induction of anti-viral CD4 and CD8 responses. The cytokine, IL-22 also displayed significant increase in young mice post infection with no change observed in adult and old mice. This is important, as IL-22 has been reported to protect mice from virus induced lung injury. No differences were detected in RANTES and IFN-γ. These results indicate that aged mice display a deficiency in production of antiviral and protective cytokines that may be responsible for their increased susceptibility to infections.

**Figure 5:**
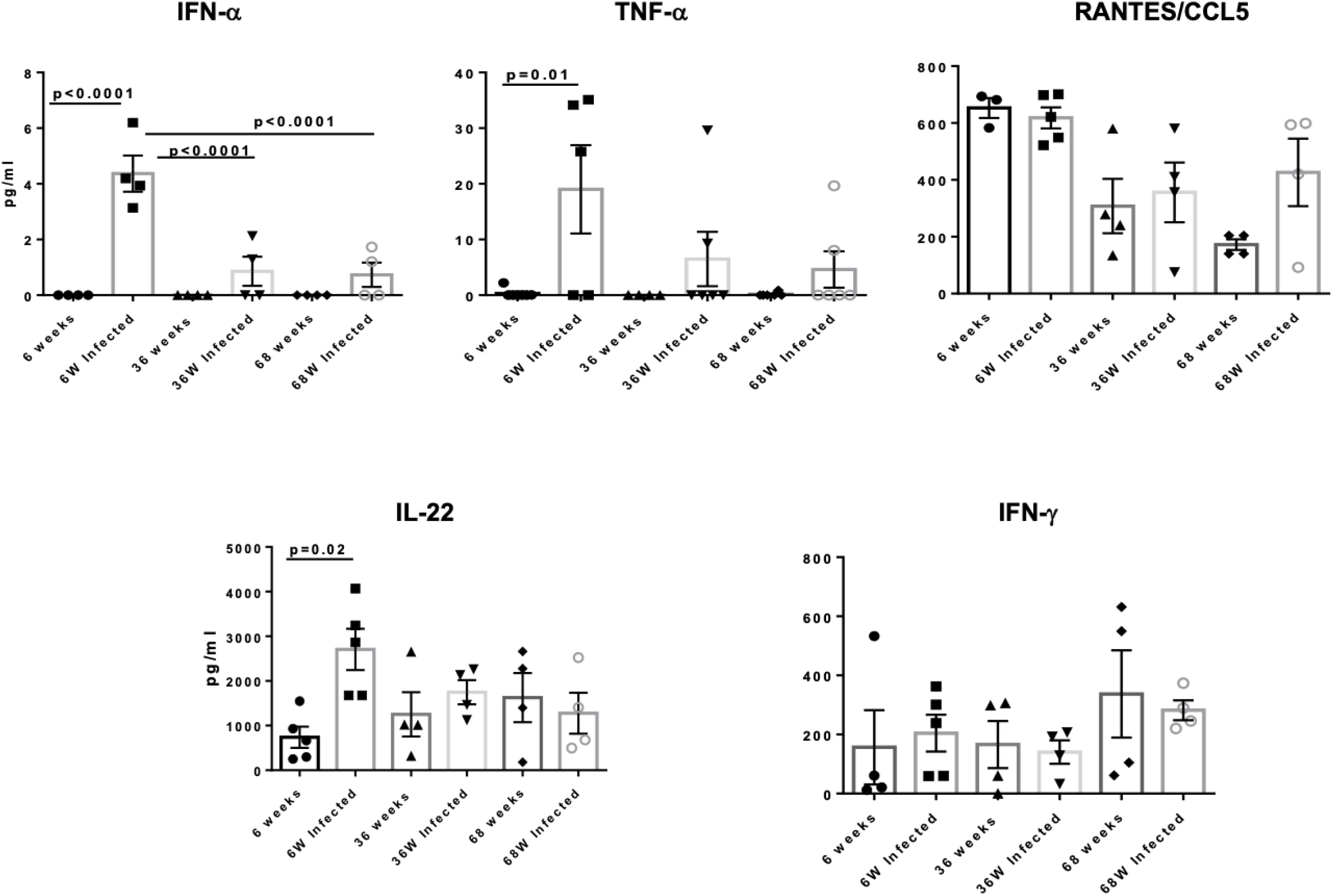
Aged mice display reduced secretion of protective cytokines in the lung after acute HSV infection. Multiplex detection was used to quantitate the levels of inflammatory mediators in the lungs of 6-week, 36-week and 68-week-old mice day 2 post infection. Graphs depict the mean +/− S.E. of the levels. N=4-5mice/group.

### 6. Viral titers and severity of HSV lung infection increases with age

Mice were inoculated intranasally with HSV-1 (KOS strain). Nasal and eye swabs and lung tissues were all collected from 6-week, 36-week and 68-week-old mice on day 2 and on day 6 for viral titer estimation. An illustration of the infection scheme and the timeline of subsequent immunological and virological assays are shown in **Fig. 6A**. The gating strategy for T cells is shown in **Fig. 6B**; lymphocytes were identified by a forward scatter (FSC) and side scatter (SSC) gate. Singlets were selected by plotting forward scatter area (FSC-A) vs. forward scatter height (FSC-H). CD4 and CD8 positive cells were then gated by the expression of CD4 and CD8. The DNA copy numbers was highest in the nasal swab and lung tissue of 68-week-old mice as compared to 36-week-old and 6-week-old mice at day 2 (**Fig. 6C**). By day 6 infection was controlled and viral titers were decreased at all age groups. In keeping with the titers, 68-week-old infected mice displayed the highest lesions followed by 6-week-old mice, while the 36-week-old-mice depicted moderate lesions (**Fig. 6D**). Altogether, this data indicates that the severity of HSV lung infection increases in aged mice.

**Figure 6:**
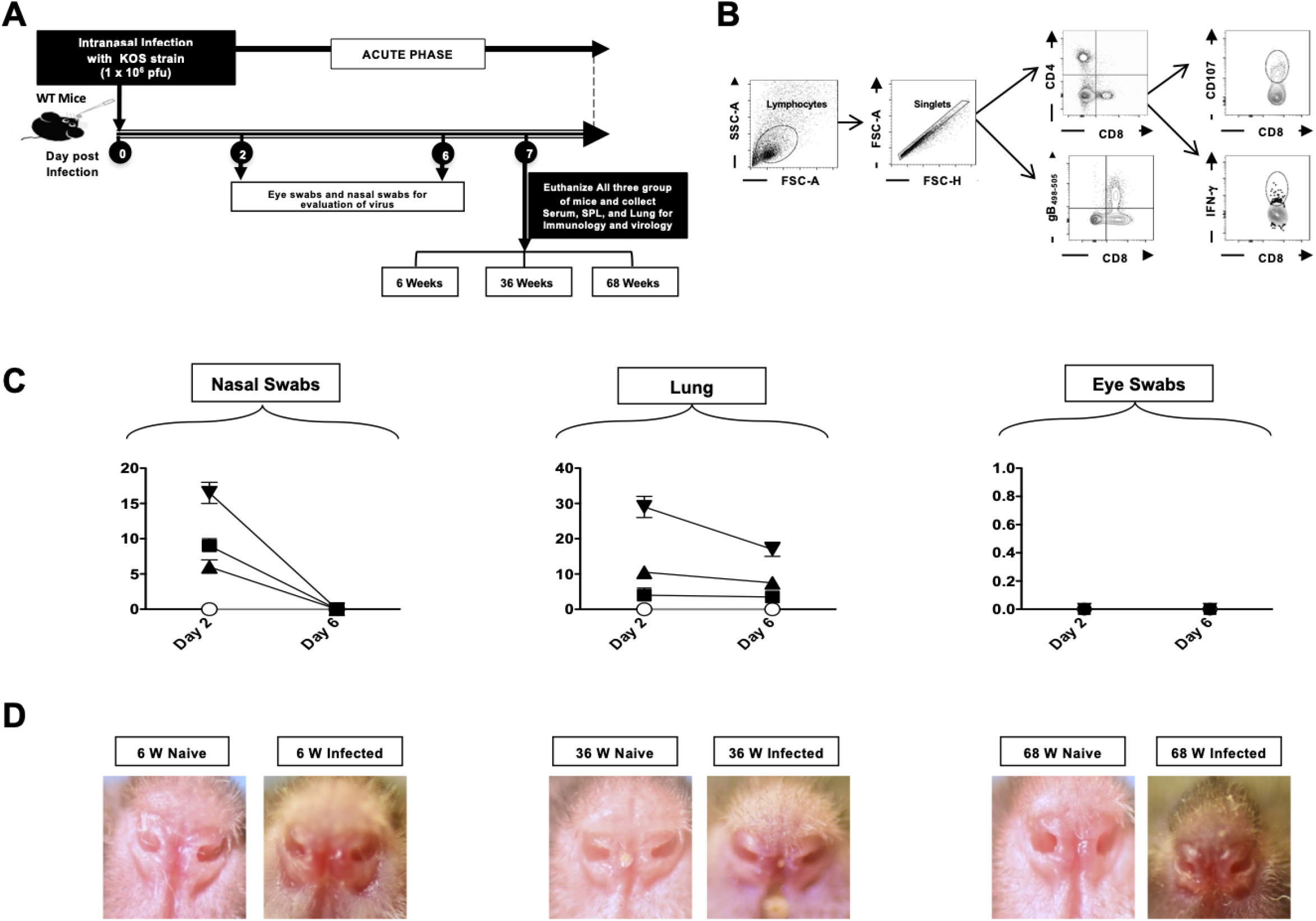
Viral titers and severity of HSV lung infection increases with age. 6-week, 36-week and 68-week-old mice were infected intranasally with 1 × 10^6^ pfu of HSV-1 strain. Mice were euthanized on day 6 post infection and single cell suspension from lungs were obtained after collagenase treatment. The lung cells were stained for T cell markers and then analyzed by FACS. (**A**) Timeline of infection and immunological analyses. (**B**) Gating strategy used to characterize lung derived cells. Lymphocytes were identified by a forward scatter (FSC) and side scatter (SSC) gate. Singlets were selected by plotting forward scatter area (FSC-A) vs. forward scatter height (FSC-H). CD8 and CD4 positive cells were then gated by the expression of CD8 and CD4 antibody. (**C**) Nasal swabs and eye swabs were collected from 6-week, 36-week and 68-week-old mice on day 2 and on day 6 for viral titer estimation. HSV-1 DNA copy numbers detected in the nasal swab, lung and eye swab on day 2 and day 6 post infection. (**D**) Representative nostril images of naïve and infected 6-week, 36-week and 68-week-old mice. Data is representative of two separate experiments.

### 7. The generation of HSV specific tetramer positive CD8^+^ T cells is impaired in aged mice

After observation of the innate immune response on day 2, we investigated the adaptive immunity in young, middle age and aged mice. On day 6 post-infection, mice were euthanized and the frequency of lung-resident CD4^+^ and CD8^+^ T cells were detected by FACS. The overall high frequencies of CD4^+^ and CD8^+^ T cells were induced in the herpes infected group as compared to the mock infected group in all three groups of mice which further declined upon aging (**Fig. 7A**). Average frequencies and absolute numbers of CD8^+^ and CD4^+^ T cells were detected in the lung of HSV-1 infected and mock-infected control group (**Fig. 7B**). Then, we compared the frequency of CD8^+^ T cells specific to HSV-1 gB_498-505_ epitope, in the lungs of 6-, 36- and 68-week old mice using the tetramers/anti-CD8 mAbs, The representative dot plots shown in **Fig. 7C** indicate increased frequencies of gB_498-505_ specific CD8^+^ T cells in the lungs of herpes infected mice which significantly decreases on aging. **Fig. 7D** shows median frequencies detected in five 6-week-old, 36-week-old and 68-week-old mice. The lowest frequency of tetramer^+^ CD8^+^ T cells was consistently detected in the aged mice as compared to the young mice (6.3% vs 11.9%), indicating that aged mice have an impaired T cell response.

**Figure 7:**
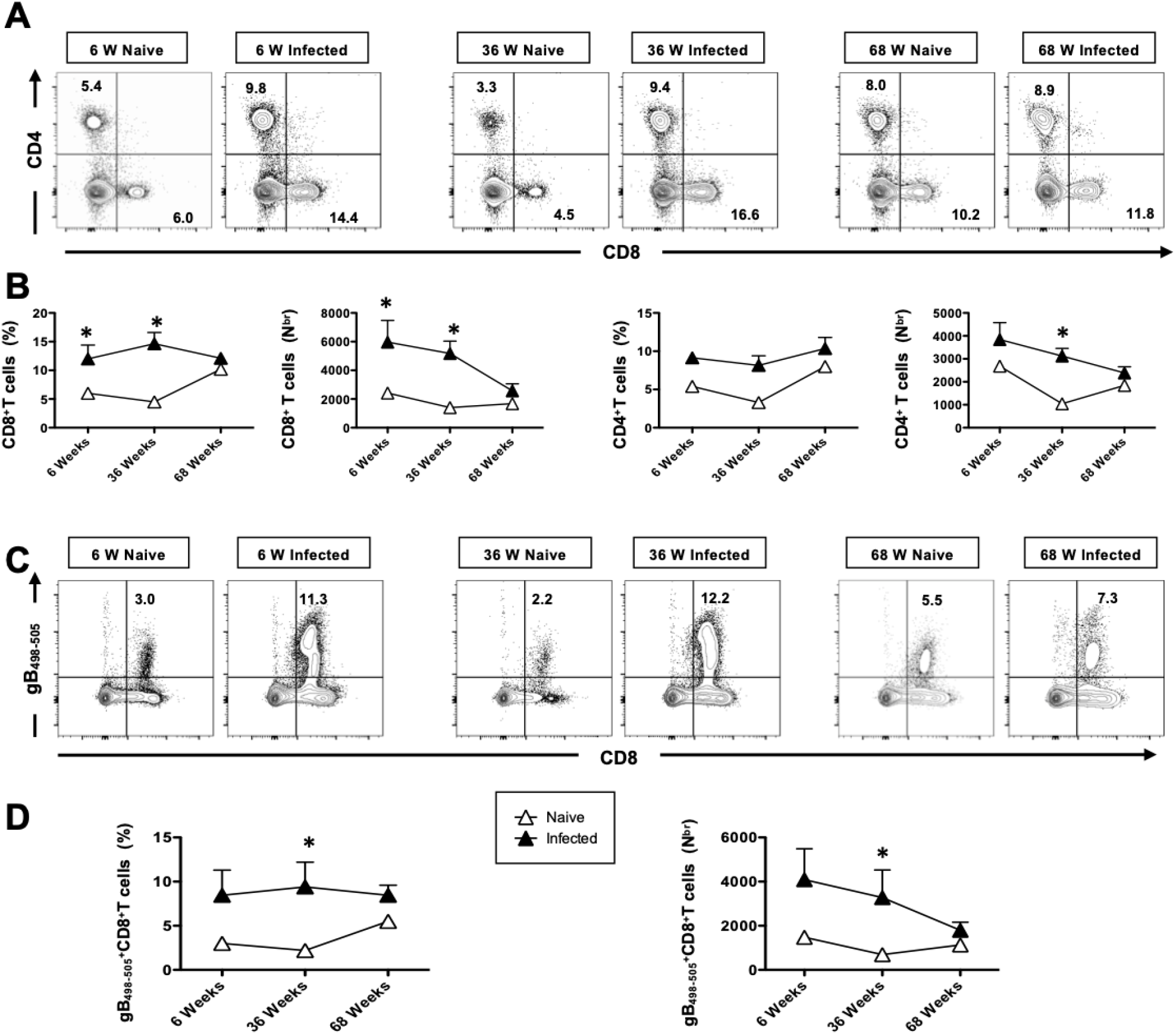
The generation of HSV specific tetramer positive CD8^+^ T cells is impaired in aged mice. 6-week, 36-week and 68-week-old mice were infected intranasally with 1 × 10^6^ pfu of HSV-1 strain. 7 days post infection mice were euthanized and single cell suspension from lungs were obtained after collagenase treatment. The lung cells were stained for T cell markers and then analyzed by FACS. (**A**) Representative FACS plots of the frequencies of CD4^+^ and CD8^+^ T cells detected in the lung of HSV-1 infected and mock-infected control group (**B**) Average frequencies and absolute numbers of CD8^+^ and CD4^+^ T cells detected in the lung of HSV-1 infected and mock-infected control group. (**C**) Representative FACS plots of the frequencies of gB_498-505_ specific CD8^+^ T cells detected in the lung of HSV-1 infected and mock-infected control group (**D**) Average frequencies and absolute numbers of gB_498-505_ specific CD8^+^ T cells detected in the lung of HSV-1 infected and mock-infected control group. Data is representative of two separate experiments.

### 8. The cytotoxic and functional activity of CD8^+^T cells is reduced with age in HSV infected mice

The impact of aging on the immune system results in defects in T cell functional responsiveness. To this end, we assessed aging-related T-cell functional response by studying CD107 and IFN-γ by T cells of young and aged mice. On day 6 post infection, mice were euthanized and a single cell suspension from the lung tissue was obtained, and the function of lung resident CD8^+^ T cells was analyzed using FACS. Significantly higher numbers of CD107^+^ CD8^+^ T cells were detected in the lungs of 6-week-old and 36-week-old HSV-1 infected mice compared to mock infected mice (**Figs. 8A** and **8B**). However, there was a significant decline in the functional response in 68-week-old mice. As shown in **Fig. 8C**, significantly high frequencies of functional IFN-γ^+^CD8^+^ T cells were detected in HSV-1 infected mice compared to mock infected mice (**Fig. 8D**) with a significant decline of these cells in the 68-week old mice. These findings indicate that HSV-1 infected 6- and 36-week-old mice induced more functional CD8^+^ T cells, however, their presence was significantly impacted with the 68-week old mice.

**Figure 8:**
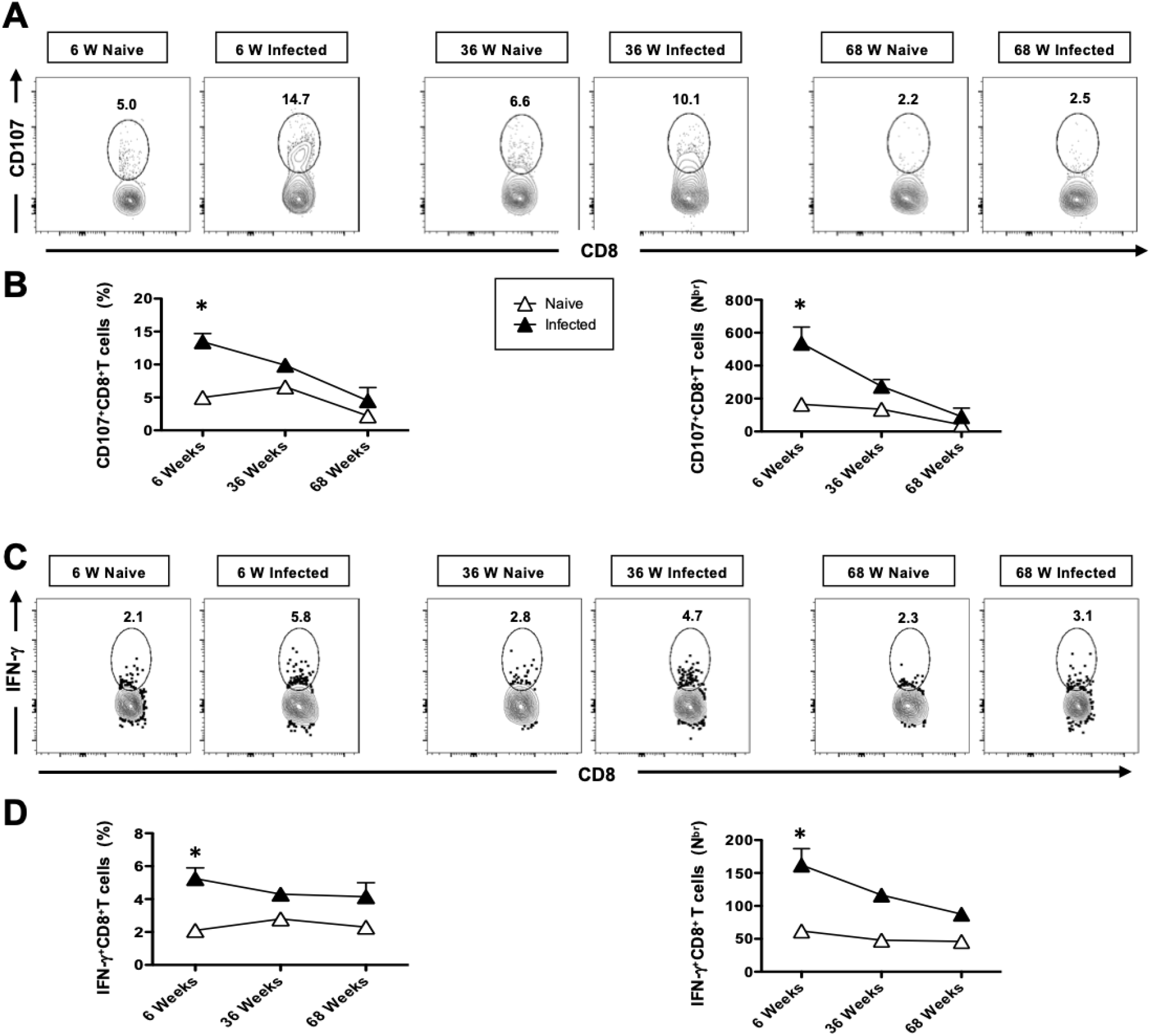
The cytotoxic and functional activity of CD8^+^T cells is reduced with age in HSV infected mice. 6-week, 36-week and 68-week-old mice were infected intranasally with 1 × 10^6^ pfu of HSV-1 strain. Seven days post infection mice were euthanized and single cell suspension from lungs were obtained after collagenase treatment. (**A**) Representative FACS plots of the frequencies of CD107^+^CD8^+^ T cells detected in the lung of HSV-1 infected and mock-infected control group (**B**) Average frequencies and absolute numbers of CD107^+^CD8^+^ T cells detected in the lung of HSV-1 infected and mock-infected control group. (**C**) Representative FACS plots of the frequencies of IFN-γ^+^CD8^+^ T cells detected in the lung of HSV-1 infected and mock-infected control group (**D**) Average frequencies and absolute numbers of IFN-γ^+^CD8^+^ T cells detected in the lung of HSV-1 infected and mock-infected control group. Data is representative of two separate experiments.

## DISCUSSION

The herpes virus affects humans of all ages; however, the elderly (>65 years old) have an increased susceptibility to these infections and are especially predisposed to severe complications (13). The increased morbidity and mortality reported in elderly populations are due to several factors that include dysfunctions in the senescent immune system.

Previously, we reported that DCs from aged subjects activate the epithelium rendering them more permeable to infections (19). Enhanced basal level inflammation/activation of cells also allows the dissemination of infections. Herein, we found that AEC functions are significantly impacted with age and, thus, may play a major role in age-associated chronic respiratory diseases and infections. AECs not only form a barrier to prevent the entry of infections but are also well-equipped to deal with, and distinguish between, innocuous and pathogenic inhalants (14). They express TLRs and other pathogen recognition receptors that allow them to sense and respond to pathogens. A decrease in ciliary beat capacity and increased secretion of mucin has been reported (15–18). In this study, we demonstrate that TLR3 expression is decreased in AECs from aged mice indicating an impairment in sensing viral nucleic acid exposed during viral replication (**Fig. 1**). This may be partially responsible for the increased incidence of other viral infections such as influenza and recently COVID-19 in the elderly.

DCs serve as the sentinels of the immune response and link innate and adaptive immunity. They are mainly comprised of two major subsets, DCs of myeloid origin and DCs of lymphoid origin. In the murine lung, different DC populations have been recently described, one of the predominant populations includes resident CD11bhigh/CD11chigh cells, (also known as conventional DCs (cDCs)). The other APC populations analyzed in these animals represent lung macrophages (CD11bhigh/CD11clow) and lymphoid DCs (CD11chigh cells) These cells have specialized antigen-processing capabilities together with co-stimulatory molecules that enable their efficient endocytosis and presentation of endogenous and exogenous antigens to initiate an immune response. DCs detect and respond to pathogens through the expression of pattern recognition receptors (PRRs) (20); (21); (22). PRRs can recognize conserved molecular components or patterns of the pathogen. Examples of PRRs include Toll-like receptors, and Nod-like receptors (20); (21). Exposure of DCs to ligands of all these PRRs results in the production of cytokines that modulate the type of T cell response and function (20); (22). Deficiencies in human TLR signaling leads to increased severity of several diseases, including sepsis, immunodeficiencies, atherosclerosis and asthma (5). We have evaluated TLR expression on cDCs, macrophages and lymphoid DC’s in the context of aging, and observed an age-associated reduced surface expression of TLR3 (**Fig. 2**). We have previously reported reduced levels of TLRs in whole blood samples and colonic biopsies from older individuals (23), and in macrophages and pDCs from aged mice (24), (25), (26). Our observation that DCs and macrophages from aged mice display reduced TLR-3 expression indicates a possible mechanism rendering the elderly being a high-risk population for viral respiratory infections including the recent COVID-19 infection.

APCs are among the first leukocytes to recognize infectious microorganisms. DCs especially are responsible for the surveillance in different tissues and subsequent migration to the lymph nodes where they interact with T-cells to present antigens and trigger the adaptive immune response. Upon encountering the antigen, APCs upregulate several molecules, such as MHC class II to present antigens to CD4^+^ T cells and provide the required second signal to fully activate these cells (27). In our study, MHC class II up-regulation was not altered in lung macrophages after herpes infection, however it was upregulated in aged DCs (myeloid and lymphoid) at the basal level, indicative of increased baseline activation of DCs that is in keeping with what has been reported for blood DCs in aging (28)(29).

Increased susceptibility to viral infection in the elderly is often ascribed to alteration in T cell functions (30–32). Contribution of the innate immune response that impacts anti-viral defense mechanisms in aging remains largely unexplored. The first line of defense after viral infections consists of robust production of interferons by innate immune cells (4, 33). These interferons have potent anti-viral properties and subsequently regulate innate and adaptive immunity (34). Numerous studies have reported a decrease secretion of IFN-α by aged pDCs, which are the most potent IFN producing cells (26, 35–37). Besides pDCs, the mDCs subset of DCs is also capable of producing a robust amount of IFNs in response to viral infection (35, 38).

In this study, we showed that IFN-α response from the lung supernatant was upregulated after herpes infection in young and adult mice, however aged mice were impaired in their capacity to secrete IFNs in response to viral infection. Our studies (37) and reports from others (26, 36, 39–41) have reported decreased production of IFN-α in the aged subjects. In addition to IFN-α, aged DCs were also deficient in the production of IL-1β and IL-22. IL-22 has emerged as a major cytokine that plays a critical role in regulating host defense and epithelial repair responses during viral infection and resolution (42). IL-22 synergizes with IL-17 to induce the secretion of antibacterial proteins and chemokines and also augments cell proliferation and repair following injury. Survival following a superinfection with *Streptococcus pneumoniae* requires IL-22 (43) attributable to protective effects of IL-22 on pulmonary epithelium. IL-22 has been shown to protect the airways by increasing transepithelial resistance and promoting bronchial epithelial cell proliferation (44). Although the IL-22R is localized to airway epithelium prior to infection, it is upregulated at parenchymal sites of lung remodeling induced by influenza (42). Furthermore, treatment of H1N1(PR8/8/34) infected mice with IL-22: Fc. caused a significant reduction in inflammation, neutrophilia, and lung leak. Importantly, this led to improved health outcomes (reduced weight loss, greater activity scores) and decreased mortality (45). IL-22 was found to prevent apoptosis through the production of anti-apoptotic proteins, such as Bcl-2 and Bcl-1, in a bleomycin model of lung injury (46). IL-22 is thus integral in protecting the host against the lung damage caused by viral infections and preventing secondary bacterial infections.

CD8 responses are highly dependent on type I IFN-induced signals, as CD8 memory formation in the setting of LCMV infection in the absence of type I IFN signaling is greatly diminished. Consistent with the delayed production of the cytokines, CD4^+^ and CD8^+^ T cells showed delayed infiltration into the lungs of aged animals (**Figs. 6A** and **6B**). In a primary infection, herpes specific cells might play a more prominent role than non-specifically activated cells. Herpes specific CD8^+^ T cells were assayed using gB_498-505_ immunodominant epitopes. Consistent with our hypothesis gB_498-505_ specific activated CD8^+^ T cells were consistently higher in young and adult mice compared to aged mice (**Figs. 7C** and **7D**). T cell responsiveness to type I IFN, commonly produced in viral and bacterial infections, is known to be pivotal for the generation of adaptive immune responses. Our findings from this report identified a reduced IFN-α response in aged mice that may contribute to reduced T cell responses. The up-regulation of the functional marker CD107 was delayed on CD8^+^ T cells in aged mice (**Fig. 8D**). Previous reports had shown no alterations of CD8^+^ T cells with age in humans and only small changes in mice (47). However, our data coincides with a recent report that demonstrated alterations in the T cell compartment of aged animals infected with influenza virus (47). T cell responsiveness to type I IFN, commonly produced in viral and bacterial infections, is known to be pivotal for the generation of adaptive immune responses. Our findings from this study identify a reduced IFN-α response in aging which may likely due to reduced expression of TLR-3. Hence, modulating TLR responses may be a beneficial immune intervention strategy for the elderly.

The aged mouse model has proven to be extremely useful in determining the effects of age on the immune responses to HSV-1 infection. In summary, our results demonstrate that HSV-infection impairs immunity in old, aged mice and propagates immune senescence. We also demonstrated that herpes infected mice exhibited perturbations in naive repertoire more profoundly than those seen in aging alone. A highly diverse T cell population is critical for protection against pathogens. The diversity of the T cell response to infection (i.e., the number of different clonotypes participating) is a better correlate of protection than the magnitude of the response (13, 41–43). As little as a 2-to 3-fold reduction in TCR repertoire diversity dramatically impairs Ag-specific responses (44, 45), and it is the T cell defects in the primary immune response that were identified as a major contributor to immune senescence. Importantly, the observed repertoire alterations were accompanied in herpes virus positive animals by altered functional responses of the remaining CD8^+^ T cells and CD4^+^ T cells and with reduced ability to clear the viral infection. We found that the absolute number of CD4^+^ and CD8^+^ T cells was lower in the older age group compared to the young group and the adult group, which indicated that the CD4^+^ and CD8^+^ T cells played important roles in controlling viral infection in aged population.

To address the functional competence of T cells in young adult and aged mice, we analyzed the production of IFN-γ. Remarkably, the increased number of polyfunctional T cells was maintained in young and adult mice but declined in aged mice. In addition, a high frequency of CD107^a^ expressing CD8^+^ T cells was also identified in the lungs of young, adult, and aged mice post infection. CD107a is a degranulation marker shown to correlate with cell-mediated cytotoxicity. Accordingly, CD3 mediated stimulation revealed that cytotoxic CD8^+^ T cells were more frequent in the lungs of young and adult mice, and their high number declined during aging. Since mature (fully primed) DCs efficiently induce cytokine production by CD8^+^ T cells and the generation of cytotoxic T cells (48), the reduced cytokine production by aged CD8^+^ T cells may be the result of reduced APC activation as suggested by alteration in MHCI and MHCII up-regulation, potentially leading to a delayed clearance of the virus.

In this study, we demonstrated that age affects the immune responses to herpes infection. Alterations in MHCII up-regulation by aged DCs and lung macrophages suggested impairments in their further activation. Remarkably, this correlated with altered levels of cytokines and chemokines, which also correlated with delayed T cell infiltration. Furthermore, herpes-specific T cells were also reduced in aged animals. These findings correlated with several reports demonstrating that age affects APC, Toll-like receptors expression and function, antigen presentation (defect in exogenous pathway), and CD8^+^ stimulating capacity (25, 49–52). However, other studies have not reported defects in DC function in the elderly (53). This might indicate that different populations of APCs at different tissues are affected differently by age. Therefore, the alterations in the APCs are most likely just one step in a large chain of alterations present in the aging immune system.

In conclusion, given the significant increase in the elderly population and increased susceptibility to herpes infection, it is becoming imperative to determine the effects of age on the immune system. Identifying the main groups of cells affected by the aging process will help to further explore the underlying mechanisms in immune alterations and design better vaccines and adjuvants to boost the immune responses in the elderly population.

## ACKNOWLEDGEMENTS

^$^Footnotes: This work is supported by Public Health Service Research R01 Grants EY026103, EY019896 and EY024618 from National Eye Institute (NEI) and R21 Grant AI158060, AI150091, AI143348, AI147499, AI143326, AI138764, AI124911 and AI110902 from National Institutes of allergy and Infectious Diseases (NIAID) (to L.BM.), and in part by The Discovery Center for Eye Research (DCER) and the Research to Prevent Blindness (RPB) grant. This work is dedicated to the memory of late Professor Steven L. Wechsler “Steve” (1948-2016), whose numerous pioneering works on herpes infection and immunity laid the foundation to this line of research.

